# Structural insight into binding site access and ligand recognition by human ABCB1

**DOI:** 10.1101/2024.08.12.607598

**Authors:** Devanshu Kurre, Phuoc X. Dang, Le T.M. Le, Varun V. Gadkari, Amer Alam

## Abstract

ABCB1 is a broad-spectrum efflux pump central to cellular drug handling and multidrug resistance in humans. However, its mechanisms of poly-specific substrate recognition and transport remain poorly resolved. Here we present cryo-EM structures of lipid embedded human ABCB1 in its apo, substrate-bound, inhibitor-bound, and nucleotide-trapped states at 3.4-3.9 Å resolution without using stabilizing antibodies or mutations and each revealing a distinct conformation. The substrate binding site is located within one half of the molecule and, in the apo state, is obstructed by transmembrane helix (TM) 4. Substrate and inhibitor binding are distinguished by major differences in TM arrangement and ligand binding chemistry, with TM4 playing a central role in all conformational transitions. Our data offer fundamental new insights into the role structural asymmetry, secondary structure breaks, and lipid interactions play in ABCB1 function and have far-reaching implications for ABCB1 inhibitor design and predicting its substrate binding profiles.

## Introduction

The ATP binding cassette (ABC) transporter ABCB1, also known as Multidrug resistance protein (MDR)1 or P-glycoprotein (p-gp) is a ubiquitously expressed drug exporter that plays a key role in cellular drug handling^1–8^. Its pharmacological relevance makes it a key transporter in the Food and Drug Administration’s guidance for all developmental drugs to be screened against^10^. ABCB1 activity can be a limiting factor in cancer chemotherapy ^6,11–14^ and treatment of neurological disorders^4,15–19^ and has been increasingly implicated in accumulation of amyloid-beta peptides, a hallmark feature of Alzheimer’s Disease^17^. Despite its relevance, ABCB1’s promise as druggable clinical target remains unrealized largely due to systemic toxicities and off target effects resulting from its inhibition^14,20^. Understanding the detailed mechanisms by which ABCB1 recognizes and transports a wide range of structurally and chemically diverse substrates remains a major focus in biomedicine. Visualizing the underlying chemistry involved is key to designing more specific ABCB1 inhibitors and circumventing ABCB1 mediated efflux of a wide range of developmental drugs. However, despite long-term efforts, ABCB1 has so far remained notoriously averse to direct structural analysis without the use of antibody fragments and stabilizing mutations to aid conformational trapping.

ABCB1 is a type II ABC exporter/type IV ABC transporter with each TMD comprising 6 transmembrane helices (TMs) and followed by a cytosolic nucleotide binding domain (NBD). It is topologically arranged as a pseudo-symmetric domain swapped dimer with the 4^th^ and 5^th^ TMs of each TMD making extensive contacts with the opposing TMDs and NBDs as first revealed by the structure of its bacterial homolog Sav1866^22^. To date, the only structures of human ABCB1 determined are those of its hydrolysis deficient mutant in the ATP bound outward facing (OF) state and those in complex with antigen binding fragments (Fabs) from the inhibitory antibodies UIC2^23^ and MRK16^24^. Key mechanistic questions about polyspecific substrate recognition and the drug transport cycle of ABCB1 therefore remain open. First, the nature of its Inward Facing (IF) apo state remains unknown, leaving open the question of show substrates gain access to their respective binding site(s). Second, the binding chemistry governing differential substrate and inhibitor interactions with ABCB1 in the absence of conformational trapping by inhibitory Fabs remains unknown. Third, it is unclear what role sequence and structural asymmetry plays in ABCB1 function. Finally, while lipids have been implicated in modulation of ABCB1 structure and its interaction with ligands ^25–28^, the extent and specifics of these interactions remains largely unexplored.

To address the above-mentioned gaps in knowledge, we determined multiple structures of wildtype human ABCB1 in a lipid environment by single particle cryo-EM. Four distinct conformations of the transporter were observed including, for the first time, its IF apo and substrate-bound states. These structures allow us to map out the conformational transitions associated with ligand and nucleotide binding and visualize key differences in how substrates and inhibitors interact with the TMD. They highlight the concerted TM and NBD movements underlying ATP coupled drug transport and regulation of binding site access and the complex interplay between lipid interactions and TM secondary structure breaks that impart tremendous TMD flexibility and overall conformational heterogeneity to human ABCB1 that has made its high-resolution structure determination difficult. Overall, our results offer fundamental insights into the mechanistic details of the ABCB1 drug transport cycle and its inhibition that will have significant implications for ABCB1 targeted therapeutic design in various medical applications as well broader drug-development efforts where potential ABCB1 interactions may limit drug-bioavailability, among other undesired effects.

### Four distinct conformations of lipid-embedded wildtype human ABCB1

Human ABCB1 (ABCB1) was stably expressed in HEK293 cells, purified in detergent, and reconstituted in saposin A (sapA) nanoparticles comprising a mixture of Brain Polar Lipids (BPL) and cholesterol (Chol). SapA reconstituted ABCB1 displayed a more homogenous mass distribution as analyzed by native mass spectrometry (nMS) as well as greater ATPase activity compared to MSP1D1 nanodisc reconstituted samples (Figure 1A-B) and was chosen for cryo-EM analysis. We analyzed ABCB1 in its apo state and in the presence of ATP/Mg^2+^ and either the substrate Taxol, representing turnover conditions similar to a recent analysis for human ABCG2^29^, or its third-generation inhibitor Zosuquidar. Taxol and Zosuquidar complexes of ABCB1 in the absence of ATP/Mg^2+^ displayed near identical conformations and are not discussed in further detail here. We also determined the structure of its nucleotide trapped state in the presence of ATP𝞬S, allowing for a visualization of the conformational spectrum associated with the drug transport cycle and its inhibition in ABCB1 (Figure 1C). The overall conformation of the zosuquidar complex was nearly identical to the inhibitor occluded state seen in the presence of UIC2 or MRK16 fabs^23,24^. Similarly, the ATP𝞬S trapped ABCB1 structure was identical to that previously reported for ATP bound state of its hydrolysis deficient EQ mutant in a detergent environment ^30^. In contrast, the conformations observed for its apo- and substrate bound states are fundamentally different and has not been previously described. Conventional models of the apo state of ABCB1 based on homologous structures or alphafold predictions invoke a symmetric, IF conformation with a wide separation between the NBDs as seen in the crystal structure of murine ABCB1^31^. Substrate binding is thought to promote NBD closure and explain consequent ATPase rate stimulation. In contrast, the apo state structure determined here displays distinct asymmetry between the two halves and closely spaced NBDs while the Taxol complex shows an IF_OPEN_ state with wider NBD separation compared to the apo conformation, among other significant differences compared to structures of the Taxol-complex of ABCB1 bound to inhibitory antibodies, as discussed in detail below.

**Figure 1.**
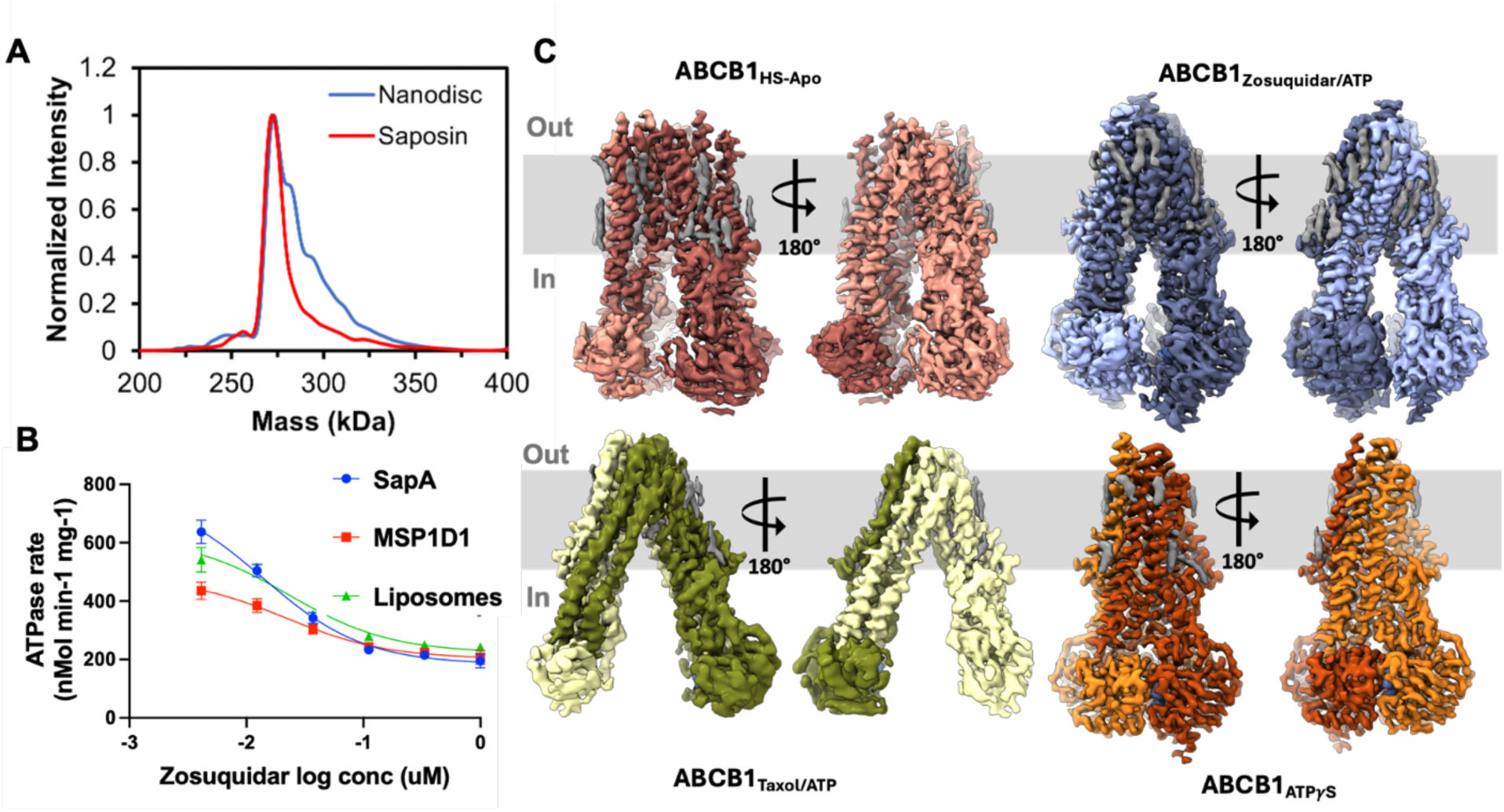
Conformational landscape of lipid embedded human ABCB1. **A C**omparison of saposin and nanodisc reconstituted human ABCB1 by nMS **B)** Comparison of ATPase activity of saposin, MSP1D1 nanodisc, and Liposome reconstituted human ABCB1. n=3 and error bars denote standard deviation. **C** Structures of human ABCB1 in multiple distinct conformational states. EM density for the two halves is colored differently and that of modeled acyl chains is colored gray.

### Apo ABCB1 adopts a unique IF_CLOSED_ conformation

The predominant conformation of apo ABCB1 observed here features an asymmetric TMD arrangement with a closed central TMD pathway (Figure 2A), closely spaced NBDs, and widely spaced extracellular “wings”^22^ (Figure 1C). We chose to classify this state as an IF_CLOSED_ state based on TMD conformation. The structure is marked by multiple secondary structure (SS) breaks in the TMDs mediated by Glycine and Proline residues and several predicted SS breakers^32^, most noticeably at G317 and G329 that leads to an elongation of extracellular loop (ECL)3 and wide separation between TM5 and TM6 (Figure S1). Conversely, ECL6, connecting TM11 & TM12 displays a lower degree of helix unraveling, likely owing to lower frequency of secondary structure breaking residues that we speculate limit its conformational freedom and possibly that of TM10 and TM11. As shown in figure 2B, closing of the central TMD pathway is facilitated by TM4, which adopts a kinked conformation with secondary structure breaks at P223 and K242, effectively dividing it into 3 sub helices (TM4a-c). In conjunction with TM6 and TM12, it forms a central 3-

**Figure 2.**
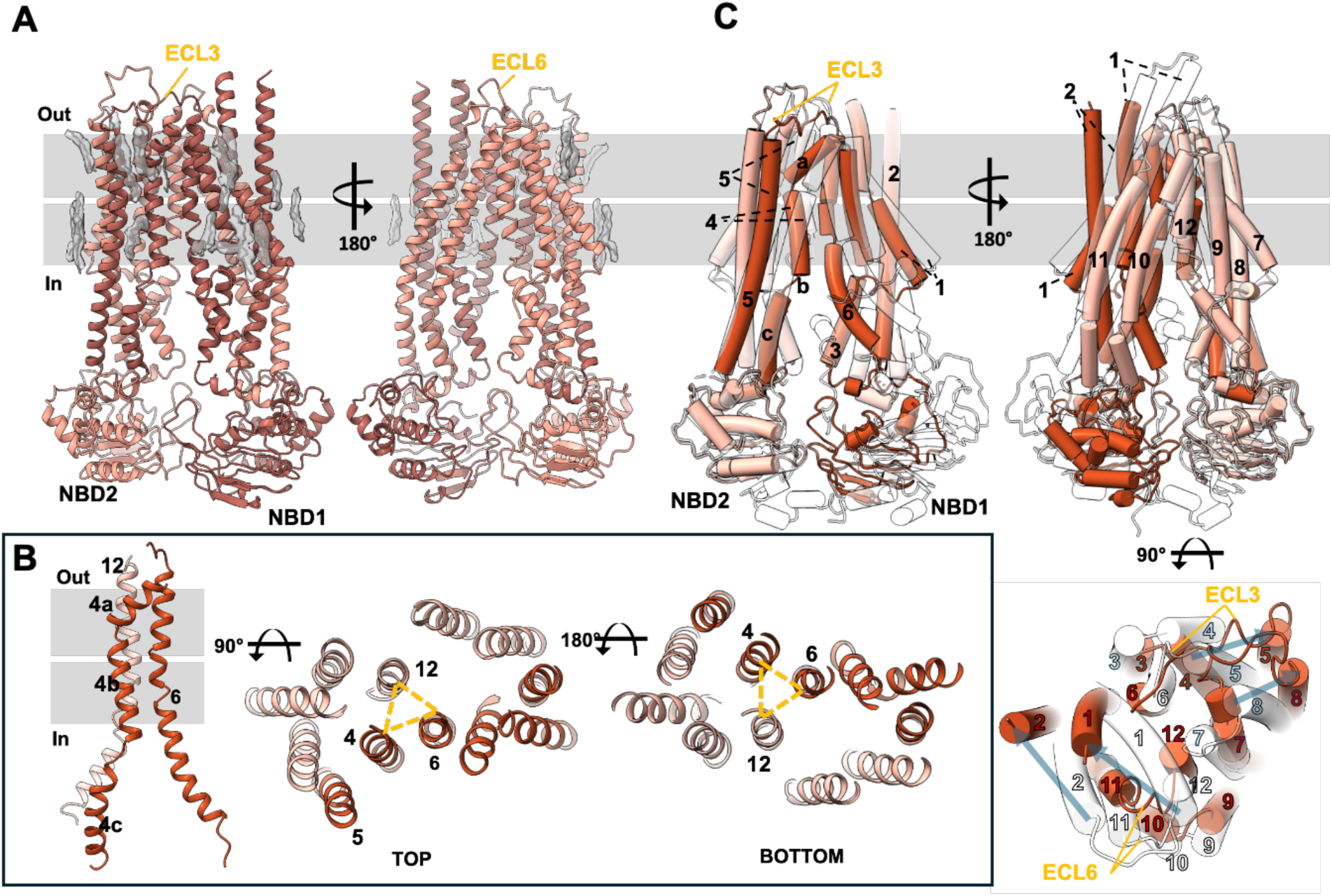
Structure of apo-ABCB1. **A** Overall structure with the two halve colored as different shades of red and density modeled as lipid acyl chains (gray sticks) shown as transparent gray surfaces. **B** 3TM bundle formation by TM4, TM6, and TM12. TM4 sub-helical segments. The yellow dashed triangle highlights the central 3TM bundle in top and bottom views. **C** Comparison of the cryo-EM structure of apo-ABCB1, colored as in A, and its alphafold predicted structure (transparent cartoon). Blue transparent arrows indicate major movements of select TMs. The gray bars represent the plasma membrane .

TM bundle that closes off the central cavity (Figure 1C). In contrast to TM4, TM10 adopts a straight conformation, contributing further to structural asymmetry and leading to a lateral opening to the lower bilayer leaflet. These features lead to an overall conformation that diverges widely from canonical IF open conformations as demonstrated by a comparison to the alphafold predicted structure of ABCB1 (Figure 2C). The starkest differences are between the respective positions of TM1/TM2 and TM4/TM5 pairs, leading to a more splayed open asymmetric arrangement of the extracellular “leaflets” and closer NBD spacing. The implications of this conformation for substrate and nucleotide access are expanded upon below.

### Distinct Substrate and Inhibitors interactions in human ABCB1

Previous analyses of substrate and inhibitor discrimination in human ABCB1 in the presence of conformational antibody Fabs revealed that both classes could occupy a centrally located, occluded TMD site with subtle differences between drug interacting residues and overall conformation^23,24^. Here we show that the predominant conformational states of Taxol and zosuquidar complexes with ABCB1 alone are completely different. As shown in Figure 3A, Taxol bound ABCB1 adopts a symmetrical IF conformation with wider NBD spacing compared to the apo state. Taxol binding, however, is asymmetric, with a single molecule observed within the C-terminal half of the molecule/2^nd^ half comprising the domain swapped (DS) TMD2 (TM7-9 and TM12 from TMD2, and TM4 and TM5 from TMD1) and NBD2 pair, offset from the central TMD space. Interestingly, this binding site is occupied by TM4b in the apo state, which swings away to allow Taxol binding (Figure 3B). This is accompanied by major rearrangements of TM5, ECL6, and TM6, breakup of the 3TM bundle observed in the apo state and an outward movement of NBD1. The position of NBD2 and its associated coupling helices remains largely unchanged. This links substrate binding to NBD orientation through TM4, which may act as an affinity gate to add a degree of substrate discrimination as expanded upon below. Density features within the hydrophobic TMD cavity are consistent with the presence of lipids and/or sterols. As their specific identity and orientation are impossible to ascertain from these data alone, we modeled them as unidentified acyl chains. A comparison of the two domain swapped halves of Taxol bound ABCB1 reveals distinct differences between residues within 5Å of the observed Taxol molecule in the C-terminal Half and its N-terminal equivalent that would present a steric and electrostatic barrier to Taxol binding. (Figure S2).

**Figure 3.**
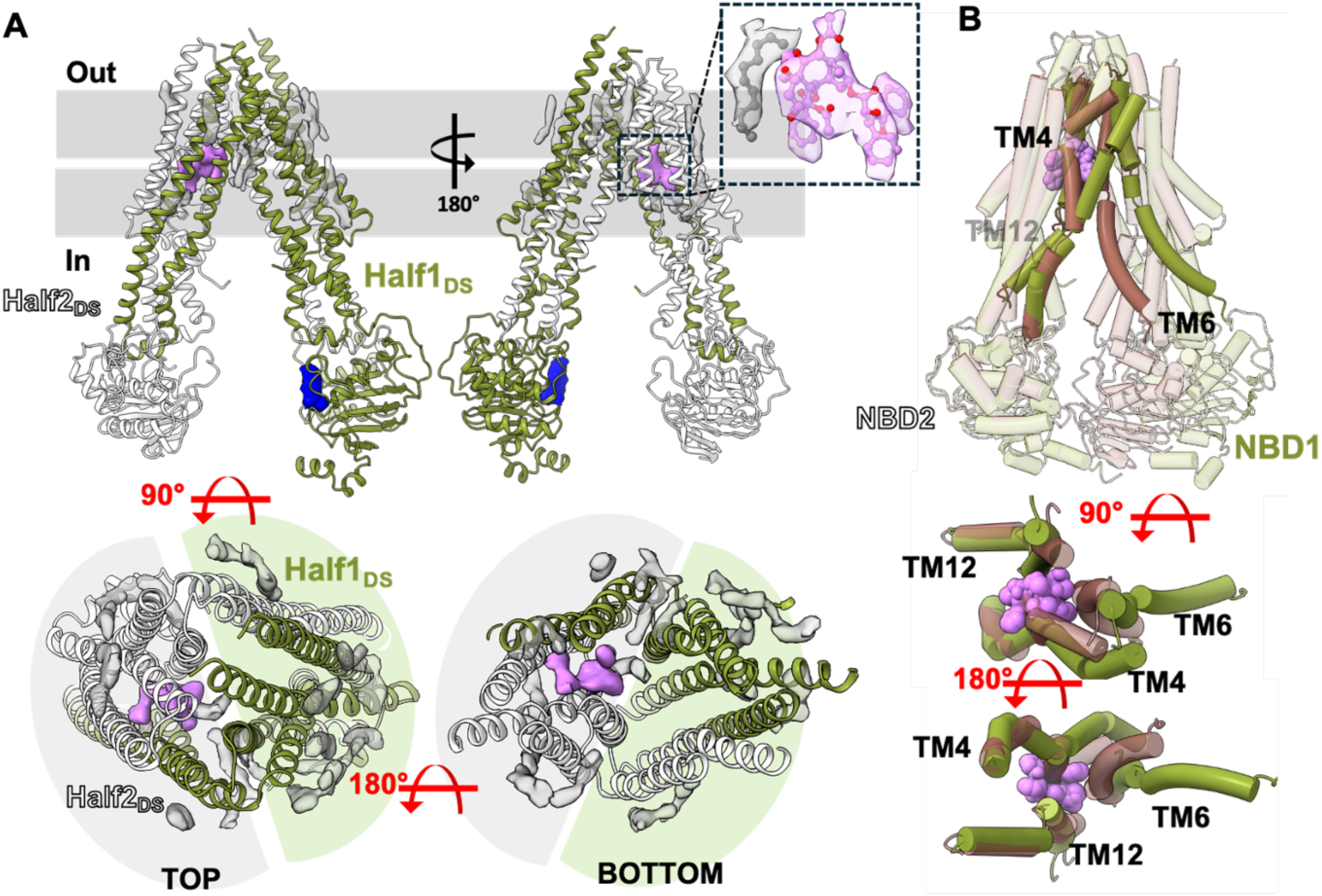
Structure of ABCB1 bound to Taxol. **A** Overall structure with first and second halves (primary structure based) colored green and white, respectively, and distinguished from domain swapped (DS) halves. Density for Taxol and lipids is shown in pink and grey (0.01 contour threshold), respectively. Weaker density for the NBD1 nucleotide is shown in blue (0.008 contour threshold). The zoom panel shows Taxol (pink sticks) density along with associated density features modeled as a lipid acyl chain (grey sticks) as transparent pink and gray surfaces, respectively. Domain swapped halves are highlighted demarcated by grey and green semicircles **B** Overall comparison of apo and Taxol complexes of ABCB1 (transparent brown and green cartoons respectively) with 3TM forming helices (solid tube helices) and Taxol (pink spheres) shown

In contrast to its Taxol complex, zosuquidar bound ABCB1 adopts the same conformation as seen in the antibody bound complexes, marked by a fully occluded cavity with 2 closely interacting zosuquidar molecules (Figure 4A-B). Cavity occlusion is brought about by the concerted kinking of TM4 and TM10, further highlighting its role in in the overall transport cycle. Diffuse density for bound nucleotide is observed in NBD1. The overall structure of zosuquidar bound ABCB1 shows increased positional order compared to the Taxol complex, with clearer density for TMD associated lipids and NBD1 associated nucleotide. While the overwhelming majority of Taxol-interacting residues are drawn from the C-terminal Half (Figure 4C), zosuquidar interactions span both halves of the transporter (Figure 4D) and no extraneous lipid density was observed in the occluded cavity.

**Figure 4.**
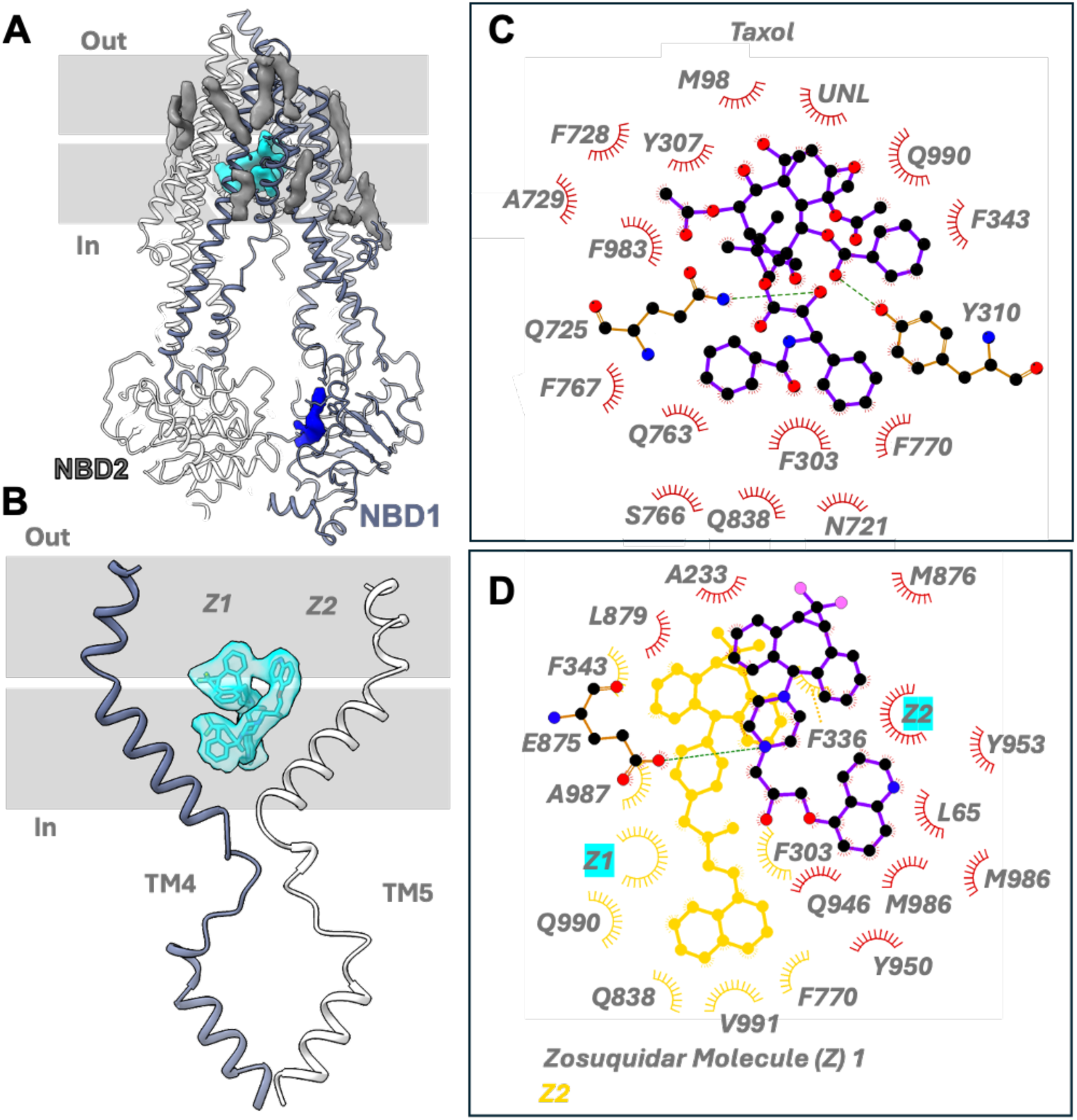
Comparison of Zosuquidar and Taxol binding. **A** Overall structure of the ABCB1 bound to Zosuquidar. Zosuquidar and ATP density is shown (0.0175 contour) as teal and blue surfaces, respectively. **B** Zoomed view of the occluded TMD cavity with TM4 and TM10 shown with EM density for both Zosuquidar molecules (teal sticks, Z1 and Z2) shown as a transparent teal surface (0.017 contour). **C** Ligand interaction plot of ABCB1 complexed to Taxol. **D** Ligand interaction plot of zosuquidar bound ABCB1 with the second Zosuquidar molecule shown in yellow.

### Structural transitions in human ABCB1 are asymmetric and dependent on TM4

The four conformational states of ABCB1 presented here allow for a direct comparison of the overall transitions associated with its drug transport cycle. As shown in Figure 5A, the C-terminal half of the transporter remains relatively rigid in comparison to its N-terminal counterpart, with significant positional changes of NBD1 associated with the different TMD conformations. Inter NBD separation is similar for the apo and inhibited state with the widest separation between the NBDs of the Taxol bound IF conformation and narrowest separation for the sandwiched NBD dimer in the ATP𝞬S complex as highlighted by Cα distance measurements between T263 (CH2) and R905 (CH4) (Figure 5B). While the overall conformations of the 4 states diverge significantly, a pairwise alignment of TM pairs 1/2, 3/6, and 4/5 (and their half 2 counterparts) shows expected patterns of linked movements during conformational cycling^33^ with major exceptions for TM4 and TM10, and to a lesser extent, TM1 and TM2 (Figure 5C). TM4 adopts a different conformation in all 4 structures, including three unique kinked conformations in the apo, substrate bound, and inhibitor bound states. Similarly, TM10 adopts different conformations in all four structures, but only the zosuquidar bound state displays a kinked conformation like that of TM4. The Cytoplasmic halves of all TMs match very closely in all structures, revealing that the conformational changes occur within the membrane environment, likely stabilized by dynamic lipid contacts, as expanded upon below.

**Figure 5.**
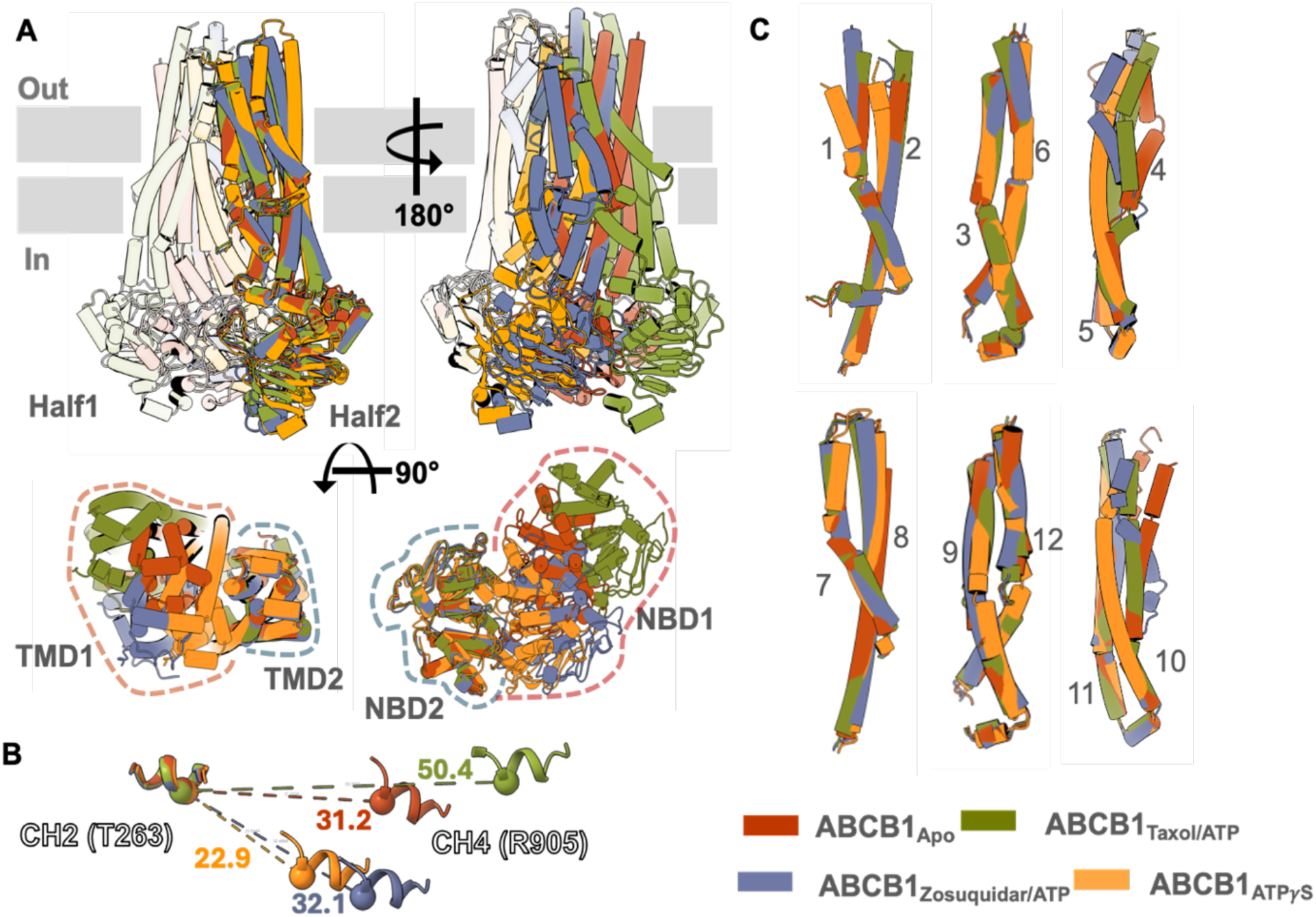
Structural Transitions in ABCB1. **A** Overlay of full transporter in all 4 conformations with Half 1 and half 2 shown as transparent surfaces (front and back views, respectively) and with individual TMDs and NBDs outlined top and bottom views, respectively. **B** Pairwise structural alignment of linked TM pairs expected to move together in different type II ABC exporter conformational states.

**Figure 6.**
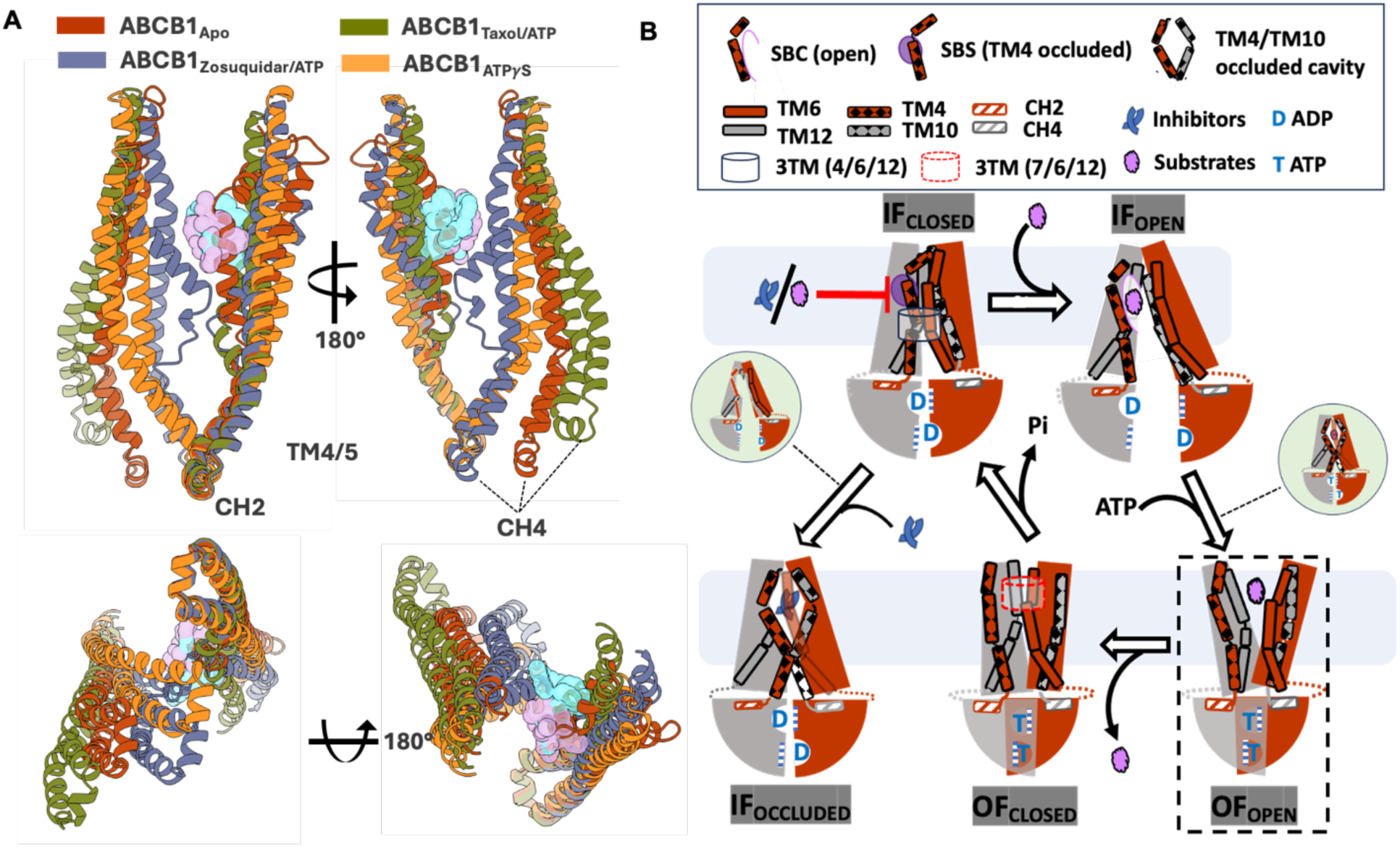
Substrate and inhibitor interactions in ABCB1. **A** Overlay of TM4/5 and TM10/11 of all structures reported, highlighting overall conformational changes linked to substrate (Taxol, pink) or inhibitor (zosuquidar, teal) binding and CH2 and CH4 movements **B** Schematic of working model for substrate transport and inhibition in human ABCB1. With the exception of the OF_OPEN_ state (based on homologous transporters like human ABCD1^9^ and Sav1866^21^). Green circles highlight potential intermediate/alternate states.

## Discussion

Our data allow us to formulate an updated mechanism for substrate transport and its inhibition in ABCB1 as shown in Figure 4B. Central to this scheme is TM4, which acts as a gating helix and undergoes large-scale rearrangements in all conformations reported here. In the unbound (apo) state, human ABCB1 likely exists in a conformational equilibrium between multiple IF states. The IF_CLOSED_ state that is dominant from our analysis is incompatible with substrate binding, with TM4 involved in 3TM bundle formation to close the TMD pathway and also sterically occluding the substrate binding site. As such, TM4 may play an autoinhibitory role and act as an affinity filter akin to the regulatory domains of ABCC type transporters^34,35^. Substrates overcoming this affinity threshold shift the conformational equilibrium towards an IF_OPEN_ state with greater NBD separation, concurrent opening of the 3-TM bundle, and ejection of TM4b from the substrate binding site. Compared to the apo state, this NBD separation may be more sterically favorable for ATP binding (ATP/ADP exchange), linking substrate binding to stimulation of ATPase rates. The IF_OCCLUDED_ state observed in the zosuquidar complex has been previously shown to be stabilized by inhibitory antibodies and capable of accommodating substrates^23,24,36^. Combined with insights from our current data, it likely represents a sparsely populated high-energy state prior to substrate extrusion through the OF_OPEN_ conformation. Inhibitors like zosuquidar, on the other hand, stabilize the IF_OCCLUDED_ conformation, thereby inhibiting the transport cycle. This clear difference between substrates and inhibitors may be explained by their divergent ligand interactions based on our observations for the Taxol and Zosuquidar complex structures described here. Asymmetry may play a key role here, with inhibitors like zosuquidar able to make stabilizing interactions with both domain-swapped halves of the transporter. In contrast, substrates like Taxol seen to bind within TMD2_DS_ may be destabilized upon contact with TMD1_DS_ upon ATP binding induced NBD closure and consequent TMD rearrangements, promoting a transition to the OF_OPEN_ conformation and substrate extrusion. This suggests that TMD1_DS_ residues that have been implicated in substrate interactions through mutagenesis and cellular efflux studies but not seen to directly interact with substrate here may be involved in promoting extrusion rather than stabilizing substrate binding^37–42^. Upon substrate extrusion the external leaflets of ABCB1 adopt a closer arrangement in contrast to OF states such as that seen in human ABCD1^9^. Interestingly, this OF_CLOSED_ state is also characterized by formation of a 3TM bundle like that in the apo state, albeit involving TM6, TM7, and TM12, and may similarly serve to prevent undesired substrate or lipid interactions before the transporter resets upon ATP hydrolysis to its IF conformation(s).

Insights into TMD access and auto-inhibition of the binding site by TM4 gleaned from our data fundamentally change our understanding of how human ABCB1 works. They have a number of important implications for development of better ABCB1 inhibitors as well as drugs that bypass its substrate transport cycle. First, the IF_CLOSED_ apo state lays the foundation for development of a new class of ABCB1 inhibitors that could potentially trap it, thereby preventing substrate access to the TMD. Second, the taxol complex offers unprecedented detail into a discreet substrate binding site that can aid the design of more accurate computational models for studying ABCB1 drug interactions. Third, the zosuquidar and taxol complexes of ABCB1 define the underlying binding chemistry that distinguishes substrates and inhibitors. Finally, our structural data reconcile decades of mutagenesis studies and showcase the remarkable structural and functional variability helix breaking elements impart to TMDs, especially in context of a lipid bilayer environment and dynamic lipid interactions, that existing homology and predicted models have failed to capture. Additional structures of human ABCB1 in complex with drugs with different physiochemical are needed to explore the extent of binding site plasticity and potential deviations from the mechanistic framework proposed above.

## Methods

### Cell Culture, Protein expression and purification

The expression and purification of wild-type human ABCB1 were conducted largely as previously described^23,24,36^. First, an ABCB1 stable cell line with a C-terminal eYFP-Rho1D4 tag and a 3C/precision protease site between the protein and tag was generated using the FIp-In TREX system (Thermo Fisher Scientific) for tetracycline-inducible expression. These ABCB1 stable cells were grown in DMEM media supplemented with 10% fetal bovine serum (FBS), penicillin-streptomycin, and antimycotic antibiotics at 37 °C in a 5% CO2 incubator until they reached over 70% confluency, which typically took about 72-96 hours. Next, the media was replaced with DMEM supplemented with 2% FBS and 0.6 µg/ml tetracycline, and the cells were allowed to express the protein for 72 hours at 37 °C in a 5% CO2 incubator. These cells were subsequently washed with PBS before being harvested by centrifugation at 3000 r.c.f for 3 minutes at 4 °C, and flash-frozen in liquid nitrogen for storage at −80 °C.

All protein purification steps were carried out at 4°C or on ice. Cell pellets were thawed and resuspended in eight volumes of lysis buffer per gram of pellet (25 mM HEPES pH 7.5, 150 mM NaCl, 20% glycerol, 0.5 mM PMSF, 2 µg/ml trypsin inhibitor, and one complete mini tablet per 50 ml). After dounce homogenizing, the cell lysate incubated with a 0.5%/0.1% mixture of n-dodecyl-β-D-maltopyranoside (DDM) and cholesteryl hemisuccinate (CHS) for 2 hours, and then centrifuged at 48000 r.c.f for 30 minutes. The supernatant was applied to Cyanogen bromide-activated Sepharose 4b beads (Cytiva) coupled to rho1D4 antibody (University of British Columbia) resin for binding over 3 hours. The resin was washed four times with 10 column volumes (CV) of wash buffer (25 mM HEPES pH 7.5, 150 mM NaCl, 20% glycerol, and 0.02%/0.004% DDM/CHS) followed by protein elution by addition of Wash Buffer supplemented either with 0.25 mg^-1^ml^-1^ 1D4 peptide (GenScript) or a 1:10 w:w ratio of 3C protease for on-column cleavage and incubated overnight at 4 °C on a roller for tag cleavage. 3C protease was removed by incubation with Ni-NTA beads.

### Lipid reconstitution of ABCB1

Expression and purification of MSP1D1 (Addgene) and Saposin A (Salipro) was carried out as described^43,44^ except that the final purification and storage buffer contained 25 mM HEPES pH 7.5, 150 mM NaCl. Brain Polar Extract lipids (BPL, Avanti) and cholesterol (Chol, Sigma) were mixed at an 80:20 w:w ratio and dried using a rotary evaporator (Bucchi), resuspended in diethyl ether, dried again, and finally resuspended in HEPES buffer (25 mM HEPES pH 7.5, 150 mM NaCl). Nanodisc reconstitution followed our published protocols^9,45^. Briefly, The BPL/Chol mixture was solubilized in storage buffer supplemented with a 0.2%/0.04% solution of DDM/CHS and homogenized using water bath sonication, with three cycles of 2 minutes on and 2 minutes off. 3C cleaved or ID4 peptide eluted ABCB1 was mixed with MSP1D1 and solubilized lipids a molar ratio of 1:10:350 for ABCB1:MSP1D1:BPL/Chol and the mixture diluted to reduce the final glycerol concentration to less than 4% (v:v). After 1-hour incubation at 4 °C on a roller, detergent was removed by addition of 0.8 grams ml^-1^ reaction buffer of Bio-Beads SM-2 (Biorad) prewashed in storage buffer and incubation on a roller for 2 hours at room temperature (RT). The supernatant was removed from the biobeads and concentrated using a 100kDA molecular weight cut off (M.W.C.O) centrifugal filter. Saposin A reconstituted ABCB1 was prepared similarly except that a 1:15:400 molar ratio of ABCB1:Saposin A:BPL/Chol was used. Protein concentration was measured by densitometry analysis of SDS-PAGE bands using detergent purified ABCB1 of known concentrations as standards.

ABCB1 proteoliposomes were prepared as described^46^ with minor modifications. Briefly, the BPL/Chol lipid mixture (80:20 wt:wt ratio) was first reconstituted in buffer comprising 150mM NaCl and 25mM Hepes pH 7.5 at a concentration of 20mg^-1^ml^-1^. Empty liposomes were prepared through extrusion using a 0.2 µm filter. Pre-formed liposomes and detergent-purified ABCB1 were supplemented with at 0.3 % and 0.14 % (v:v) of Triton X-100, respectively, mixed, and incubated at RT for 30 minutes. Detergent removal was done in five successive incubation steps using each using fresh 50 mg Bio-beads SM-2 per ml reaction mix. The incubation steps were carried out with gentle agitation for 30 mins at RT, 60 mins at 4°C, overnight at 4 °C, followed by two 60-minute incubations at 4 °C. Liposomes were pelleted by ultracentrifugation at 80,000 r.p.m using a TLA-100 rotor (Beckmann Coulter), the supernatant discarded and resuspended in an equivalent volume of reconstitution buffer at 0.5-1 mg^-1^ml^-1^.

### ATPase Assays

ATPase measurements were based on a molybdate-based calorimetric assay measuring release of inorganic phosphate (Pi) ^47^ as described^9,45^. Stocks of zosuquidar (Tocris) and taxol (PhytoLab) were prepared in 100% Dimethyl sulfoxide DMSO. ATPase measurements were performed by incubating 0.02-0.1 mg^-1^ml^-1^ ABCB1 with 2mM ATP, 10mM MgCl_2_ with varying concentrations of zosuquidar or taxol at 37°C. Statistical analyses and linear regression were done in GraphPad Prism 9. All assays were replicates of three independent experiments.

### Native-Mass spectrometry

Wild-type ABCB1 was purified and reconstituted into nanoparticles as described in the above sections. The detergent sample and the reconstituted ABCB1 samples were buffer exchanged into 200 mM ammonium acetate (99.999% Trace Metals Basis, Sigma Aldrich) containing 0.02% DDM/0.004% CHS (only 200 mM ammonium acetate for nanoparticle sample) using 40k zeba spin desalting column and further purified by injecting into an Agilent 1260 Infinity II LC system (Agilent Technologies) using pre-equlibrated TSKgel G4000SWxl column (TOSOH biosciences) Samples were diluted to 500 nM and ionized via nano-electrospray ionization using gold coated borosilicate capillaries (prepared in-house) and analyzed on a Q Exactive Ultra High Mass Range orbitrap mass spectrometer (Thermo Fisher Scientific)^48,49^. The instrument was operated in Direct Mass mode, enabling orbitrap-based charge detection mass spectrometry measurements of individual intact lipoprotein nanoparticle ions^50,51^. Briefly, the instrument was operated with the Ion Target set to “high m/z” and the Detector Optimization set to “low m/z.” The in-source trapping and higher-energy collisional dissociation cell were operated at 1-10 V. All measurements were acquired at a resolution setting of 200,000 (FWHM at m/z 400) with a trapping gas pressure setting of 1. All data processing was performed using STORIBoard (Proteinaceous Inc.). Ions were filtered based on ion lifetime and signal-to-noise, and ion charge states were assigned using the “Voting v3” charge assignment algorithm^51^. Ion filtering and charge assignment parameters are summarized in Table S1. Charge assignment was calibrated using carbonic anhydrase, alcohol dehydrogenase, pyruvate kinase, beta-galactosidase, and GroEl. All samples were acquired for 10-20 minutes, and the reported measurements are representative of ∼10,000 ions.

### Cryo EM Sample Preparation & Data collection

For Grid preparation ABCB1-eYFP reconstituted in Saposin A Nanoparticles (SapNPs) were incubated antiGFP nanobody (Addgene) coupled Sepharose 4B resin prepared in house for 2 hrs at 4°C, washed with 3 x 10CV of reconstitution buffer, followed on-column cleavage by in 3CV reconstitution buffer supplemented with 3C protease to recover ABCB1 SapNPs. Samples were subsequently concentrated using a 100 MWCO centrifugal filter and further purified by Size exclusion chromatography (SEC) on an Agilent 1260 Infinity II LC system (Agilent Technologies) using a TSKgel G4000SWxl column (TOSOH biosciences) pre-equilibrated with reconstitution buffer at 4 °C and peak fractions pooled and concentrated to 0.5-1 mg^-1^ml^-1^ for grid preparation . Where needed zosuquidar and taxol were added to pooled fractions at 10 µM final concentration with or without ATP/Mg^2+^ (5mM each) and incubated for 10 minutes at RT before concentration. 4 µL of sample was applied to the glow discharged (60 s, 15 mA) Quantifoil R1.2/1.3 Cu grids using Vitrobot Mark IV with 4 s blot time and 0 blot force under >90 % humidity at 4 °C and plunge frozen in liquid ethane. All grids were clipped and stored in liquid nitrogen.

All the Cryo EM data were collected on a 300 kV Titan Krios electron microscope equipped with a Biocontinuum K3 Direct Electron Detector with 20 eV GIF energy filter, 50 eV condenser C2 and 100 µm objective apertures. Automated data collection was carried out using the EPU 2.8.0.1256REL software package (Thermo Fisher Scientific) at a magnification of 130,000× in Counted Super Resolution mode corresponding to a calibrated pixel size of 0.664 Å with defocus range set from −0.5 µm to −2.5 µm. Three shots were taken per hole. Image stacks comprising 40 frames were recorded for 60 s at an estimated dose rate of 1e-/Å^2^/frame.

### Data processing, model building, and refinement

Data processing was done in Relion^52–54^. In brief, image stacks were motion corrected using Relion’s internal MotionCor2 implementation, followed by CTF estimation using CTFFIND4^55^. All resolution estimates were based on the gold standard 0.143 cutoff criterion^53^. Data collection and processing parameters are provided in Table S2 along with model building and refinement statistics. Data processing flow charts are shown in Figure S3. EM density around individual domains/TMs and Local resolution-colored maps are shown in Figure S5 and Figure S6, respectively.

For ABCB1-apo, an initial dataset comprising 5974 micrographs was used for reference free automated particle picking (Laplacian-of-Gaussian algorithm) within Relion. 2167202 particles were extracted at a 3x binned pixel size of 1.992 Å and subjected to several rounds of 2D classification, followed by Ab-initio model building using within Relion. This initial model was used for subsequent 3D classification (number of classes (N)=5) and a single predominant class comprising 662694 was refined to 5.1 Å followed by another round of 3D classification (N=5) and 3D refinement, re-extraction at a 1.5X binned pixel size of 0.996 Å, and particle polishing to yield a 4.1 Å map. A second set of 6321903 particles from 13327 micrographs was picked using Topaz (default model) and processed similarly except that a refined 3D class from the first set was used as a reference. A refined 3D at 4.0 Å resolution and comprising 660276 particles was obtained. Particles from the final refined classes from both sets were combined, followed by additional rounds of 3D refinement and postprocessing to yield a 3.8 Å map.

For the ABCB1_Taxol/ATP_ complex, 15494460 particles from 33055 micrographs were autopicked using Topaz and extracted at a 3x binned pixel size of 1.992 Å. After one round of 2D Classification, 6254156 particles were used for 3D classification (N=5) with a low pass filtered ABCB1-apo map as a reference. The single highest resolution class revealed an IF conformation and was subjected to iterative 3D refinement and particle polishing, followed by subtraction of the SapNP. After 3D classification (N=5), 154538 particles from the highest resolution were reverted to their original non-subtracted images and refined to 3.9 Å.

For the ABCB1_Taxol_ complex, 5725 micrographs were used to pick 2547172 particles by Topaz and extracted at a 3x binned pixel size of 1.992 Å. After 2D classification, 1270596 particles were used for 3D classification (N=3) using the ABCB1_apo_ map as a reference. 486111 particles from the best class were subjected to another round of 3D classification. The single highest resolution class comprised 133895 particles and was refined to 4.7 Å.

For the ABCB1_Zosuquidar_ complex, 2182930 particles were automatically picked by Topaz from 7281 micrographs. After two rounds of 2D classification, 943398 particles entered 3D classification (N=5) with a low-pass filtered ABCB1_apo_ map used as a reference The single, highest resolution class comprising 373279 particles was subjected to re-extraction at a 1.5X binned pixel size of 0.996 Å and signal subtraction to remove delocalized bulk lipid density and refined to 3.6 Å resolution.

For the ABCB1_Zosuquidar/ATP_ complex, 10710935 particles from 12897 micrographs were picked using topaz. 2468729 particles were chosen for 3D classification (N=5) using the map of the zosuquidar complex without ATP as a reference. A single highest resolution class comprising 733688 particles was subjected to iterative rounds of 3D classification and particle polishing within Relion to yield a final refined map at 3.6 Å resolution.

Fort the ABCB1_ATP𝞬S_ sample, 7689616 particles from 12165 micrographs were automatically picked using Topaz. After several rounds of 2D classification, 1732065 particles were subjected to 3D classification (N=5)using a low pass filtered ABCAB1_Apo_ map as a reference. A single OF classes comprising 400787 particles was subjected to another round of 3D classification (N=5). 180,163 Particles from two similar and roughly equally populated OF classes were combined, re-extracted at a 1.5X binned pixel size of 0.996 Å and refined to 3.75 Å. A second dataset of 6204620 particles from 9318 micrographs was processed similarly to yield a final refined class at 3.5 Å comprising 260172 particles. Particles from the final class from both datasets were combined and subjected to another round of 3D classification (N=5) and the highest resolution class comprising 136896 particles was refined to 3.4 Å.

Final EM maps were used for model building in COOT 0.9.6 EL^56^. De novo model building was guided by the predicted structure of ABCB1 from AlphaFold2^57^ for the apo and taxol complexes. For the zosuquidar complexes, model building was guided initially by the structure of ABCB1 bound to the MRK16 fab (PDBID 7A6F). For the ATP𝞬S complexed ABCB1, the structure of ATP bound ABCB1-EQ (PDBID: 6C0V) was used as an initial model before minor adjustments and refinement. Non-proteinaceous continuous density features attributed to lipids or sterols were modeled as Acyl-chains. The structures were refined with secondary structure and geometry restrains in COOT 0.9.6 and PHENIX^58^. Where NBD density was too weak for denovo model building, docked NBDs from higher resolution structures reported here were used and minimally refined. The final models for ABCB1_apo_ comprised residues 33-81, 106-606, 694-1230, for ABCB1_Taxol/ATP_ comprised residues 30-87, 100-630, 689-1257, for ABCB1_Zosuquidar/ATP_ comprised residues 30-90, 104-630, 691-1272, and for ABCB1_ATP𝞬S_ comprised residues 35-80, 105-630, 692-1276. Map and Structure visualization was performed in UCSF Chimera^59^ and ChimeraX^60^.

## Acknowledgments

We would like to thank Dr. Kaspar Locher at ETH, Zurich, Switzerland, for providing the synthetic gene construct of human ABCB1. We would also like to thank the cryo-EM and shared instruments core facilities at the Hormel Institute for help with experimental setup, and Dr. Jeppe Olsen, Dr. Jarrod French, Dr. Thanuja Sudasinghe, Dr. Subhrajyoti Dola, and Ashley Wise for critical reading and discussion during manuscript preparation. This work was supported in part by the Hormel Foundation (Institutional research funds to AA), the National Institutes of Health (NIH) 1R01GM146906 (to AA), the Eagles Telethon postdoctoral fellowship (LTML and DK). V.V.G acknowledges funding from University of Minnesota start-up funds.

## Author Contributions

AA conceived the research. DK performed all experiments with contributions from LTML, AA, and PXD. AA and DK performed all cryo-EM Data processing, model building, and refinement. V.V.G performed the nMS data collection analysis simulations. AA and DK wrote the manuscript with contributions from all other authors.

## Data and materials availability

Requests for materials should be addressed to Amer Alam. The cryo-EM Maps have been deposited at the Electron Microscopy Databank (EMDB) under accession codes EMD-45854 (ABCB1_apo_), EMD-45904 (ABCB1_Taxol/ATP_), EMD-45903 (ABCB1_Zosuquidar/ATP_), and EMD-45906 (ABCB1_ATP𝞬S_) and the associated atomic coordinates have been deposited at the Protein Data bank (PDB) under accession codes 9CR8, 9CTF, 9CTC, and 9CTG, respectively. Maps for ABCB1_Taxol_ and ABCB1_Zosuquidar_ have been deposited at the EMDB with accession codes MD-45931 and EMD-45932, respectively.

## Additional Information

Supplementary Information is available for this manuscript.

## Supplementary Data

**Table S1:** Ion filtering and charge assignment parameters.

| **Ion Filtering** |  |
| --- | --- |
| R<sup>2</sup> Threshold | 0.996 |
| Duration threshold | 0.42 |
| Minimum Time of Death | 0.2 |
| Maximum Time of Birth | 0.1 |
| Signal-To-Noise Threshold | 3 |
| **Charge Assignment (Voting v3)** |  |
| Bin Size (ppm) | 5 |
| Minimum Ions in Bin | 1 |
| Number of Charge Neighbors | 2 |
| Number of Isotope Neighbors | 5 |

**Table S2.** Cryo-EM data collection, refinement and validation statistics of human ABCB1.

| **Data collection and processing** | ABCB1<sub>apo</sub> | ABCB1<br>Taxol/ATP | ABCB1<br>Zosuquidar/ATP | ABCB1<br>ATP<sub>γ</sub>S | ABCB1<br>Zosuquidar | ABCB1<br>Taxol |
| --- | --- | --- | --- | --- | --- | --- |
| Microscope | FEI Titan Krios |  |  |  |  |  |
| Camera | Gatan Biocontinuum K3 |  |  |  |  |  |
| Magnification | 130kx |  |  |  |  |  |
| Voltage (kV) | 300 |  |  |  |  |  |
| Electron exposure (e<sup>-</sup>/Å<sup>2</sup>) | 45-50 |  |  |  |  |  |
| Defocus range (μm) | -0.5 to -2.5 |  |  |  |  |  |
| Pixel size (Å) | 0.664 |  |  |  |  |  |
| Energy filter (eV) | 20 |  |  |  |  |  |
| Micrographs (#) |  |  |  | 9318 | 7281 |  |
| Symmetry imposed | C1 |  |  |  |  |  |
| Particles in final Class | 660276 | 154538 | 733688 | 136896 | 373279 | 133895 |
| Map resolution (Å)<br>(FSC 0.143) | 3.8 | 3.9 | 3.6 | 3.4 | 3.6 | 4.7 |
| Map sharpening B factor (Å<sup>2</sup>) | -197.74 | -<br>168.435 | -223.113 | -137.7 | -<br>148.229 | -<br>240.975 |
| Local Resolution Range (Å<sup>2</sup>) | 3.7-4.9 | 3.8-5.9 | 3.5-4.5 | 3.4-5.0 |  |  |
| **Refinement** | ABCB1<sub>apo</sub> | ABCB1<br>Taxol/ATP | ABCB1<br>Zosuquidar/ATP | ABCB1<sub>ATP<sub>γ</sub>S</sub> |  |  |
| Model composition |  |  |  |  |  |  |
| Non-hydrogen atoms | 8664 | 9398 | 9525 | 9128 |  |  |
| Protein residues |  |  |  |  |  |  |
| Ligands | 1087<br>15 | 1158<br>24 | 1170<br>25 | 1157<br>13 |  |  |
| B factors (Å<sup>2</sup>) |  |  |  |  |  |  |
| Protein | 65.21 | 128.74 | 79.80 | 70.65 |  |  |
| Ligand | 41.21 | 100.83 | 60.54 | 62.68 |  |  |
| R.m.s. deviations |  |  |  |  |  |  |
| Bond lengths (Å) | 0.003 | 0.002 | 0.003 | 0.004 |  |  |
| Bond angles (°) | 0.489 | 0.483 | 0.516 | 0.579 |  |  |
| Validation |  |  |  |  |  |  |
| MolProbity score | 1.84 | 1.62 | 1.73 | 1.59 |  |  |
| Clashscore | 7.50 | 9.52 | 6.72 | 7.68 |  |  |
| Poor rotamers (%) | 0.22 | 0.52 | 0.31 | 0.11 |  |  |
| Ramachandran plot |  |  |  |  |  |  |
| Favored (%) | 93.52 | 97.40 | 94.85 | 97.05 |  |  |
| Allowed (%) | 6.48 | 2.52 | 4.81 | 2.95 |  |  |
| Disallowed (%) | 0.0 | 0.09 | 0.34 | 0 |  |  |

**Figure S1.**
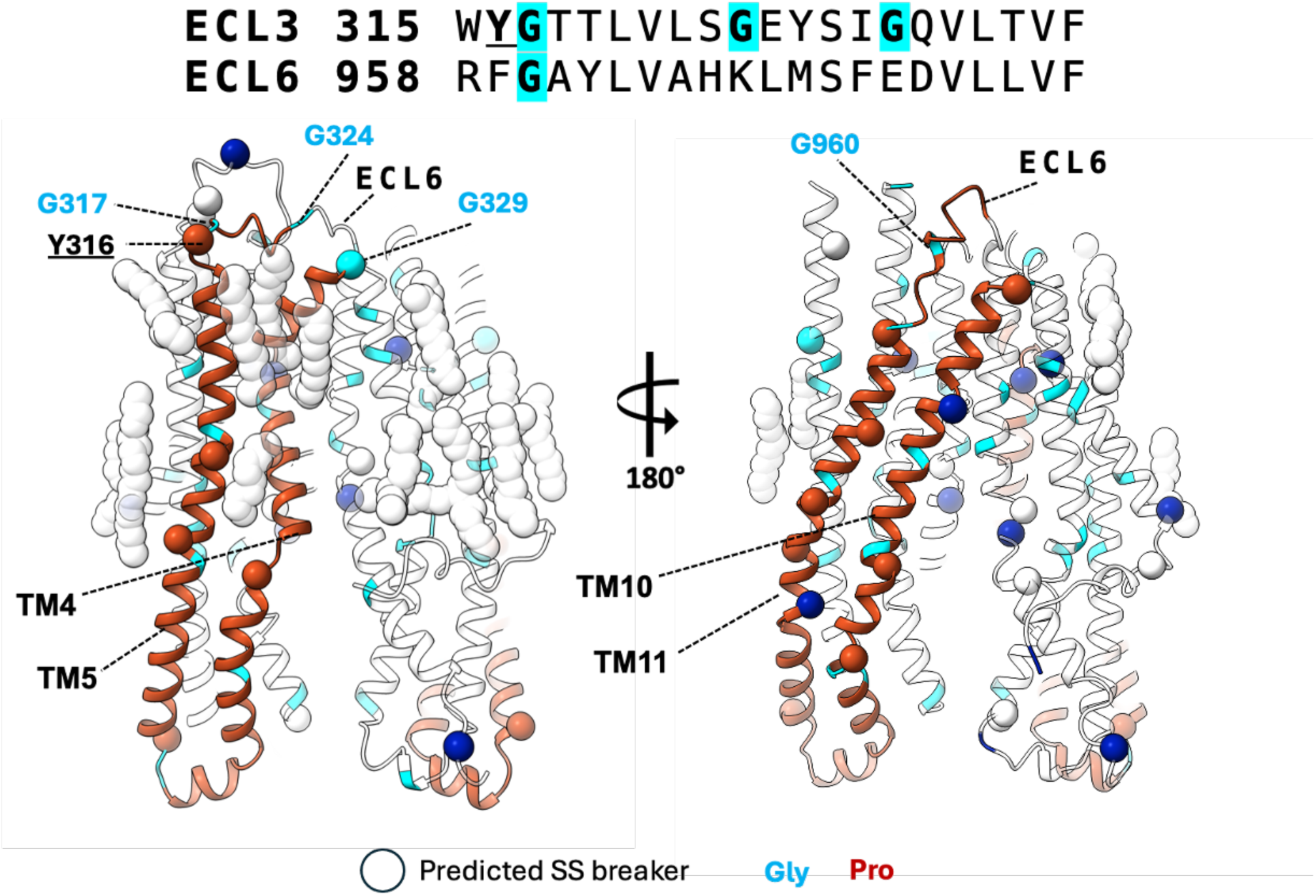
Secondary structure (SS) breaks in apo ABCB1. Gly and Pro residues colored teal and blue, respectively, and predicted SS breaks shown as spheres. An ECL3 and ECL6 sequence alignment is also shown with residues colored similarly and predicted SS breaking residues underlined. TM4/5 and TM10/11 pairs are colored red. Acyl chains for prospective lipid/sterol molecules are shown as transparent spheres.

**Figure S2.**
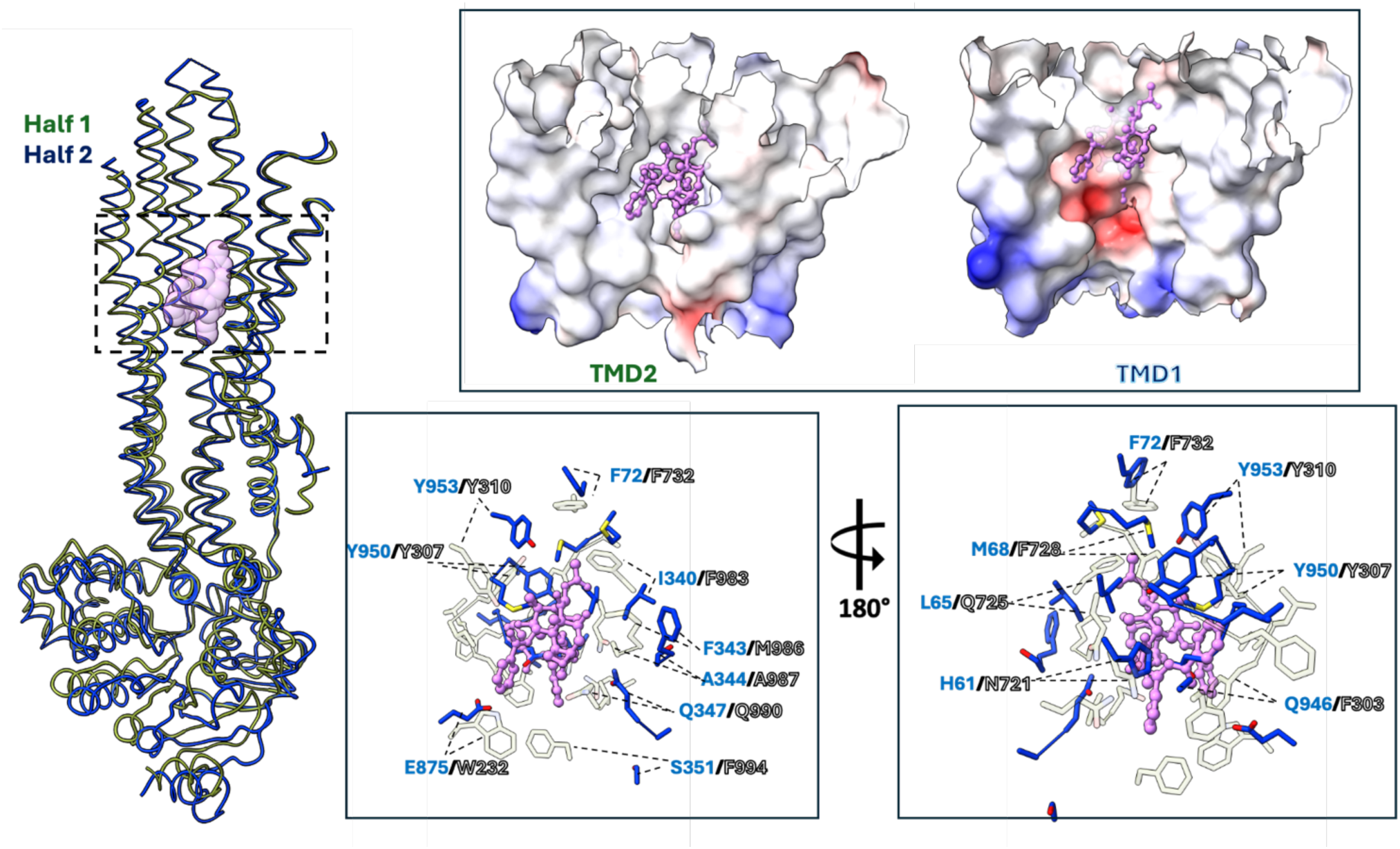
Mismatch between TMD1 and TMD2 cavities for taxol binding. **A** Overlay of domain swapped (DS) halves of ABCB1. The Taxol molecule bound to TMD2DS is shown as transparent pink spheres. The Zoom panel shows electrostatic potential map of the TMD2DS cavity (left) and its TMD1DS cavity equivalent (right) showing electrostatic and steric clashes with Taxol. **B** TMD1DS equivalent residues of TMD2DS residues (Blue sticks) within 5 Angstroms of bound Taxol (transparent sticks), with residue labels colored similarly.

**Figure S3.**
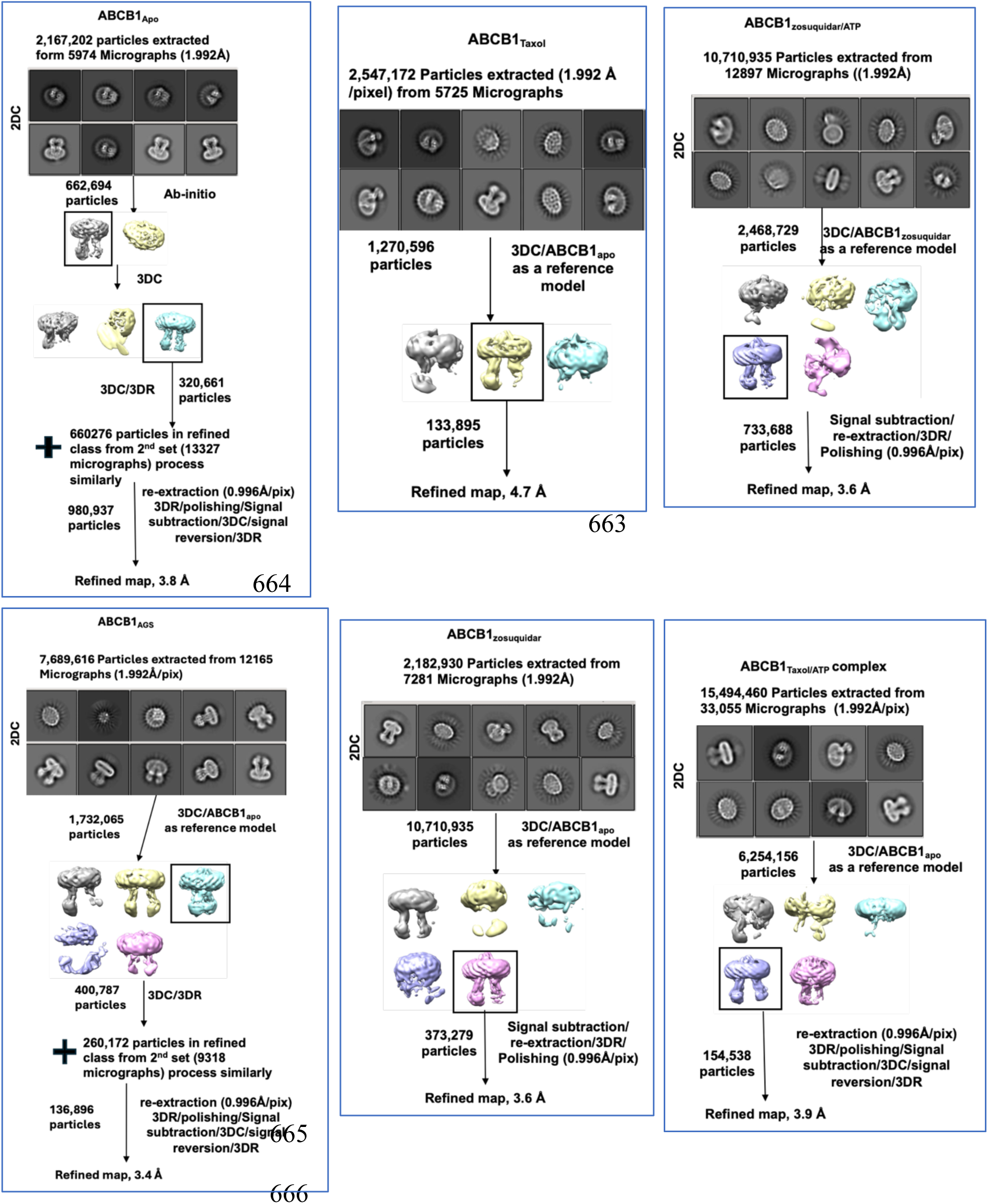
Data processing overview. 3D classes chosen for further processing are boxed.

**Figure S4.**
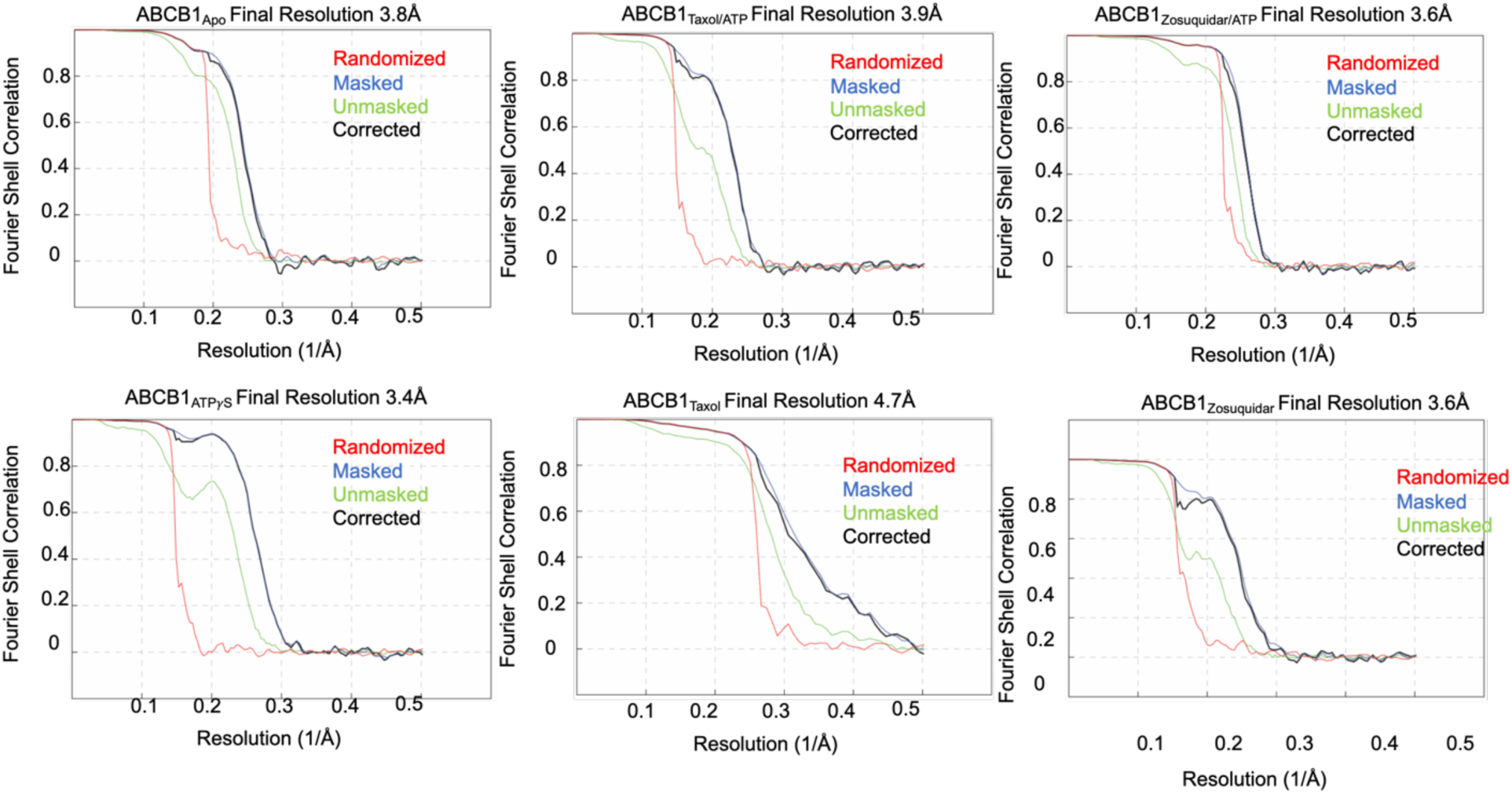
Data processing overview. FSC curves for final human ABCB1 maps

**Figure S5.**
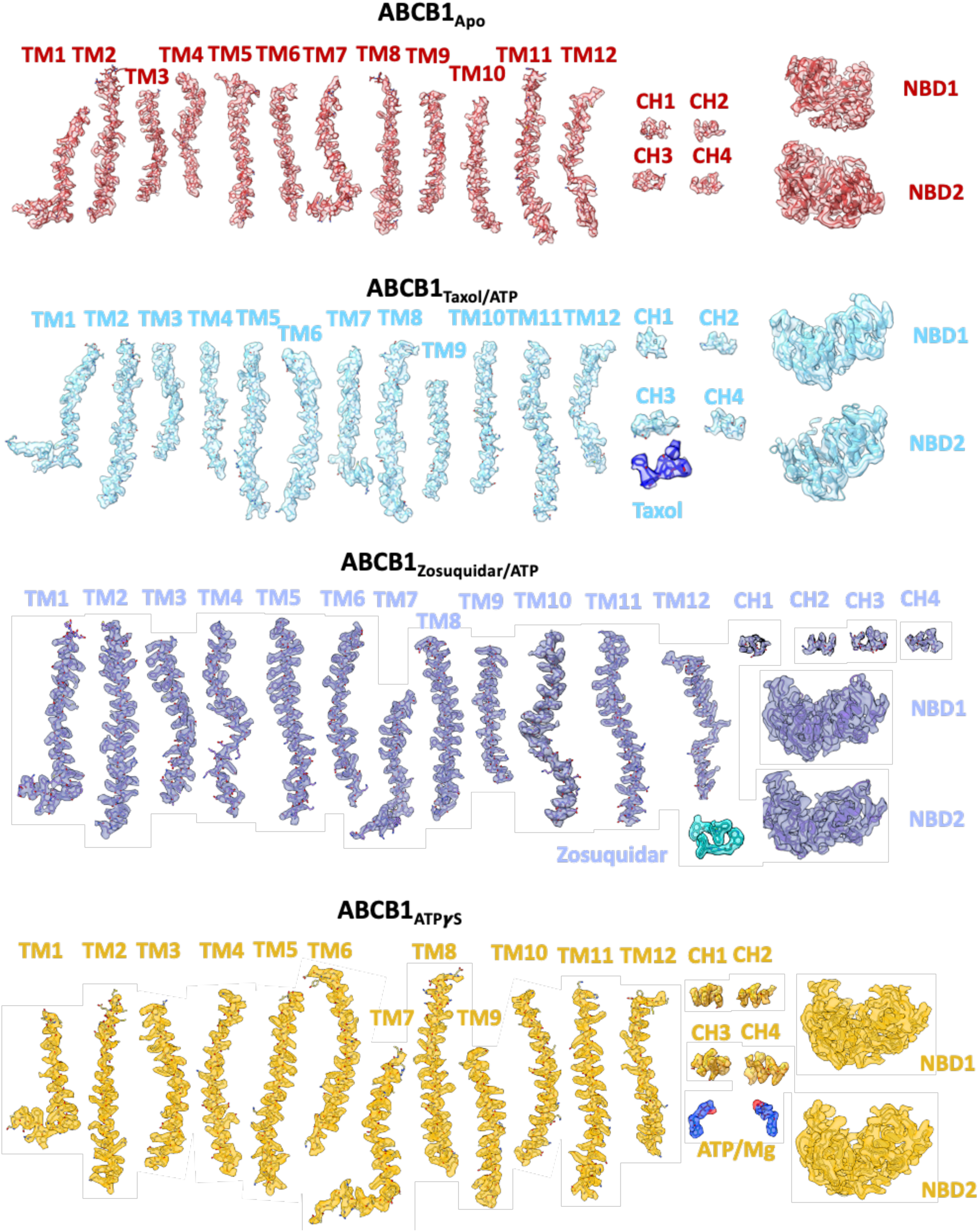
EM Density maps for lipid embedded ABCB1. Contour levels for Apo: 0.012; Taxol/ATP complex: 0.011; Zosuquidar/ATP complex: 0.031; and ATP𝞬S complex:0.011

**Figure S6.**
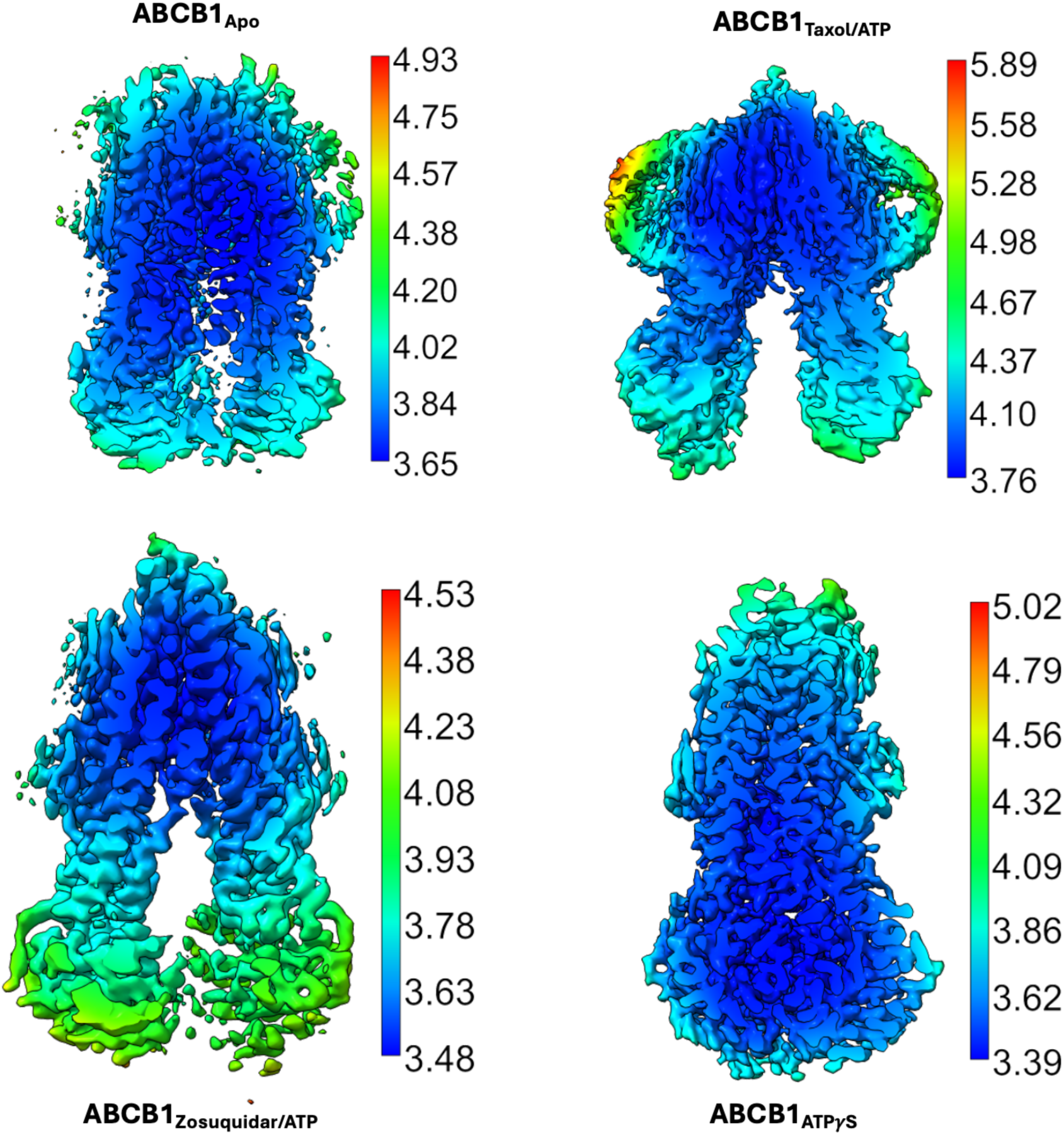
Local Density filtered Maps. Color Key indicates Resolution range for each filtered map

## References

1 Borst, P. & Schinkel, A. H. P-glycoprotein ABCB1: a major player in drug handling by mammals. J Clin Invest 123, 4131–4133, doi:10.1172/JCI70430 (2013).

2 Hodges, L. M. et al. Very important pharmacogene summary: ABCB1 (MDR1, P-glycoprotein). Pharmacogenet Genomics 21, 152–161, doi:10.1097/FPC.0b013e3283385a1c (2011).

3 Leslie, E. M., Deeley, R. G. & Cole, S. P. Multidrug resistance proteins: role of P-glycoprotein, MRP1, MRP2, and BCRP (ABCG2) in tissue defense. Toxicol Appl Pharmacol 204, 216–237, doi:10.1016/j.taap.2004.10.012 (2005).

4 Fromm, M. F. Importance of P-glycoprotein at blood-tissue barriers. Trends Pharmacol Sci 25, 423–429, doi:10.1016/j.tips.2004.06.002 (2004).

5 Borst, P. & Elferink, R. O. Mammalian ABC transporters in health and disease. Annu Rev Biochem 71, 537–592, doi:10.1146/annurev.biochem.71.102301.093055 (2002).

6 Ueda, K., Cardarelli, C., Gottesman, M. M. & Pastan, I. Expression of a full-length cDNA for the human “MDR1” gene confers resistance to colchicine, doxorubicin, and vinblastine. Proc Natl Acad Sci U S A 84, 3004–3008, doi:10.1073/pnas.84.9.3004 (1987).

7 Thiebaut, F. et al. Cellular localization of the multidrug-resistance gene product P-glycoprotein in normal human tissues. Proc Natl Acad Sci U S A 84, 7735–7738, doi:10.1073/pnas.84.21.7735 (1987).

8 Darwich, A. S., Neuhoff, S., Jamei, M. & Rostami-Hodjegan, A. Interplay of metabolism and transport in determining oral drug absorption and gut wall metabolism: a simulation assessment using the “Advanced Dissolution, Absorption, Metabolism (ADAM)” model. Curr Drug Metab 11, 716–729 (2010).

9 Le, L. T. M., Thompson, J. R., Dang, P. X., Bhandari, J. & Alam, A. Structures of the human peroxisomal fatty acid transporter ABCD1 in a lipid environment. Commun Biol 5, 7, doi:10.1038/s42003-021-02970-w (2022).

10 Fiedorczuk, K. & Chen, J. Mechanism of CFTR correction by type I folding correctors. Cell 185, 158–168 e111, doi:10.1016/j.cell.2021.12.009 (2022).

11 Bauer, F. et al. Synthesis and in vivo evaluation of [11C]tariquidar, a positron emission tomography radiotracer based on a third-generation P-glycoprotein inhibitor. Bioorg Med Chem 18, 5489–5497, doi:10.1016/j.bmc.2010.06.057 (2010).

12 Leonard, G. D., Fojo, T. & Bates, S. E. The role of ABC transporters in clinical practice. Oncologist 8, 411–424, doi:10.1634/theoncologist.8-5-411 (2003).

13 Ling, V. Multidrug resistance: molecular mechanisms and clinical relevance. Cancer Chemother Pharmacol 40 **Suppl**, S3–8 (1997).

14 Robey, R. W. et al. Revisiting the role of ABC transporters in multidrug-resistant cancer. Nat Rev Cancer 18, 452–464, doi:10.1038/s41568-018-0005-8 (2018).

15 Schinkel, A. H., Wagenaar, E., Mol, C. A. & van Deemter, L. P-glycoprotein in the blood-brain barrier of mice influences the brain penetration and pharmacological activity of many drugs. J Clin Invest 97, 2517–2524, doi:10.1172/JCI118699 (1996).

16 Xie, R., Hammarlund-Udenaes, M., de Boer, A. G. & de Lange, E. C. The role of P-glycoprotein in blood-brain barrier transport of morphine: transcortical microdialysis studies in mdr1a (-/-) and mdr1a (+/+) mice. Br J Pharmacol 128, 563–568, doi:10.1038/sj.bjp.0702804 (1999).

17 Storck, S. E., Hartz, A. M. S. & Pietrzik, C. U. The Blood-Brain Barrier in Alzheimer’s Disease. Handb Exp Pharmacol 273, 247–266, doi:10.1007/164_2020_418 (2022).

18 Sita, G., Hrelia, P., Tarozzi, A. & Morroni, F. P-glycoprotein (ABCB1) and Oxidative Stress: Focus on Alzheimer’s Disease. Oxid Med Cell Longev 2017, 7905486, doi:10.1155/2017/7905486 (2017).

19 Loscher, W. & Potschka, H. Role of multidrug transporters in pharmacoresistance to antiepileptic drugs. J Pharmacol Exp Ther 301, 7–14, doi:10.1124/jpet.301.1.7 (2002).

20 Tamaki, A., Ierano, C., Szakacs, G., Robey, R. W. & Bates, S. E. The controversial role of ABC transporters in clinical oncology. Essays Biochem 50, 209–232, doi:10.1042/bse0500209 (2011).

21 Dawson, R. J. & Locher, K. P. Structure of the multidrug ABC transporter Sav1866 from Staphylococcus aureus in complex with AMP-PNP. FEBS Lett 581, 935–938, doi:10.1016/j.febslet.2007.01.073 (2007).

22 Dawson, R. J. & Locher, K. P. Structure of a bacterial multidrug ABC transporter. Nature 443, 180–185, doi:10.1038/nature05155 (2006).

23 Alam, A., Kowal, J., Broude, E., Roninson, I. & Locher, K. P. Structural insight into substrate and inhibitor discrimination by human P-glycoprotein. Science 363, 753–756, doi:10.1126/science.aav7102 (2019).

24 Nosol, K. et al. Cryo-EM structures reveal distinct mechanisms of inhibition of the human multidrug transporter ABCB1. Proc Natl Acad Sci U S A 117, 26245–26253, doi:10.1073/pnas.2010264117 (2020).

25 Clay, A. T., Lu, P. & Sharom, F. J. Interaction of the P-Glycoprotein Multidrug Transporter with Sterols. Biochemistry 54, 6586–6597, doi:10.1021/acs.biochem.5b00904 (2015).

26 Hegedus, C., Telbisz, A., Hegedus, T., Sarkadi, B. & Ozvegy-Laczka, C. Lipid regulation of the ABCB1 and ABCG2 multidrug transporters. Adv Cancer Res 125, 97–137, doi:10.1016/bs.acr.2014.10.004 (2015).

27 Loo, T. W. & Clarke, D. M. P-glycoprotein ATPase activity requires lipids to activate a switch at the first transmission interface. Biochem Biophys Res Commun 472, 379–383, doi:10.1016/j.bbrc.2016.02.124 (2016).

28 Szewczyk, P. et al. Snapshots of ligand entry, malleable binding and induced helical movement in P-glycoprotein. Acta Crystallogr D Biol Crystallogr 71, 732–741, doi:10.1107/S1399004715000978 (2015).

29 Yu, Q. et al. Structures of ABCG2 under turnover conditions reveal a key step in the drug transport mechanism. Nat Commun 12, 4376, doi:10.1038/s41467-021-24651-2 (2021).

30 Kim, Y. & Chen, J. Molecular structure of human P-glycoprotein in the ATP-bound, outward-facing conformation. Science 359, 915–919, doi:10.1126/science.aar7389 (2018).

31 Aller, S. G. et al. Structure of P-glycoprotein reveals a molecular basis for poly-specific drug binding. Science 323, 1718–1722, doi:10.1126/science.1168750 (2009).

32 Imai, K. & Mitaku, S. Mechanisms of secondary structure breakers in soluble proteins. Biophysics (Nagoya-shi) 1, 55–65, doi:10.2142/biophysics.1.55 (2005).

33 Lee, J. Y., Yang, J. G., Zhitnitsky, D., Lewinson, O. & Rees, D. C. Structural basis for heavy metal detoxification by an Atm1-type ABC exporter. Science 343, 1133–1136, doi:10.1126/science.1246489 (2014).

34 Mao, Y. X. et al. Transport mechanism of human bilirubin transporter ABCC2 tuned by the inter-module regulatory domain. Nat Commun 15, 1061, doi:10.1038/s41467-024-45337-5 (2024).

35 Khandelwal, N. K. & Tomasiak, T. M. Structural basis for autoinhibition by the dephosphorylated regulatory domain of Ycf1. Nat Commun 15, 2389, doi:10.1038/s41467-024-46722-w (2024).

36 Alam, A. et al. Structure of a zosuquidar and UIC2-bound human-mouse chimeric ABCB1. Proc Natl Acad Sci U S A 115, E1973–E1982, doi:10.1073/pnas.1717044115 (2018).

37 Loo, T. W. & Clarke, D. M. Mapping the Binding Site of the Inhibitor Tariquidar That Stabilizes the First Transmembrane Domain of P-glycoprotein. J Biol Chem 290, 29389–29401, doi:10.1074/jbc.M115.695171 (2015).

38 Loo, T. W. & Clarke, D. M. Mutational analysis of ABC proteins. Arch Biochem Biophys 476, 51–64, doi:10.1016/j.abb.2008.02.025 (2008).

39 Loo, T. W. & Clarke, D. M. Mutational analysis of human P-glycoprotein. Methods Enzymol 292, 480–492 (1998).

40 Chufan, E. E., Kapoor, K. & Ambudkar, S. V. Drug-protein hydrogen bonds govern the inhibition of the ATP hydrolysis of the multidrug transporter P-glycoprotein. Biochem Pharmacol 101, 40–53, doi:10.1016/j.bcp.2015.12.007 (2016).

41 Chufan, E. E. et al. Multiple transport-active binding sites are available for a single substrate on human P-glycoprotein (ABCB1). PLoS One 8, e82463, doi:10.1371/journal.pone.0082463 (2013).

42 Nasim, F. et al. Active transport of rhodamine 123 by the human multidrug transporter P-glycoprotein involves two independent outer gates. Pharmacol Res Perspect 8, e00572, doi:10.1002/prp2.572 (2020).

## References (Methods)

43 Ritchie, T. K. et al. Chapter 11 - Reconstitution of membrane proteins in phospholipid bilayer nanodiscs. Methods Enzymol 464, 211–231, doi:10.1016/S0076-6879(09)64011-8 (2009).

44 Frauenfeld, J. et al. A saposin-lipoprotein nanoparticle system for membrane proteins. Nat Methods 13, 345–351, doi:10.1038/nmeth.3801 (2016).

45 Le, L. T. M. et al. Cryo-EM structures of human ABCA7 provide insights into its phospholipid translocation mechanisms. EMBO J 42, e111065, doi:10.15252/embj.2022111065 (2023).

46 Geertsma, E. R., Nik Mahmood, N. A., Schuurman-Wolters, G. K. & Poolman, B. Membrane reconstitution of ABC transporters and assays of translocator function. Nat Protoc 3, 256–266, doi:10.1038/nprot.2007.519 (2008).

47 Chifflet, S., Torriglia, A., Chiesa, R. & Tolosa, S. A method for the determination of inorganic phosphate in the presence of labile organic phosphate and high concentrations of protein: application to lens ATPases. Anal Biochem 168, 1–4 (1988).

48 Wilm, M. & Mann, M. Analytical properties of the nanoelectrospray ion source. Anal Chem 68, 1–8, doi:10.1021/ac9509519 (1996).

49 Fort, K. L. et al. Expanding the structural analysis capabilities on an Orbitrap-based mass spectrometer for large macromolecular complexes. Analyst 143, 100–105, doi:10.1039/c7an01629h (2017).

50 Worner, T. P. et al. Resolving heterogeneous macromolecular assemblies by Orbitrap-based single-particle charge detection mass spectrometry. Nat Methods 17, 395–398, doi:10.1038/s41592-020-0770-7 (2020).

51 Kafader, J. O. et al. STORI Plots Enable Accurate Tracking of Individual Ion Signals. J Am Soc Mass Spectrom 30, 2200–2203, doi:10.1007/s13361-019-02309-0 (2019).

52 Zivanov, J. et al. A Bayesian approach to single-particle electron cryo-tomography in RELION-4.0. Elife 11, doi:10.7554/eLife.83724 (2022).

53 Scheres, S. H. RELION: implementation of a Bayesian approach to cryo-EM structure determination. J Struct Biol 180, 519–530, doi:10.1016/j.jsb.2012.09.006 (2012).

54 Kimanius, D. et al. Data-driven regularization lowers the size barrier of cryo-EM structure determination. Nat Methods, doi:10.1038/s41592-024-02304-8 (2024).

55 Rohou, A. & Grigorieff, N. CTFFIND4: Fast and accurate defocus estimation from electron micrographs. J Struct Biol 192, 216–221, doi:10.1016/j.jsb.2015.08.008 (2015).

56 Brown, A. et al. Tools for macromolecular model building and refinement into electron cryo-microscopy reconstructions. Acta Crystallogr D Biol Crystallogr 71, 136–153, doi:10.1107/S1399004714021683 (2015).

57 Jumper, J. et al. Highly accurate protein structure prediction with AlphaFold. Nature 596, 583–589, doi:10.1038/s41586-021-03819-2 (2021).

58 Adams, P. D. et al. PHENIX: a comprehensive Python-based system for macromolecular structure solution. Acta Crystallogr D Biol Crystallogr 66, 213–221, doi:10.1107/S0907444909052925 (2010).

59 Pettersen, E. F. et al. UCSF Chimera--a visualization system for exploratory research and analysis. J Comput Chem 25, 1605–1612, doi:10.1002/jcc.20084 (2004).

60 Pettersen, E. F. et al. UCSF ChimeraX: Structure visualization for researchers, educators, and developers. Protein Sci 30, 70–82, doi:10.1002/pro.3943 (2021).

